# Patient-Derived Circulating Monocytes Promote Calcific Aortic Valve Disease Progression

**DOI:** 10.64898/2026.04.30.721898

**Authors:** Léa Di Maria, Hugues Boël, Nicolas Perzo, Sylvanie Renet, Chloé Valentin, Théo Lemarcis, Bryan Marais, Zina Badji, Thomas Levesque, Delphine Béziau-Gasnier, Hélène Eltchaninoff, Ebba Brakenhielm, Eric Durand, Sylvain Fraineau

## Abstract

**Background:** Calcific aortic valve disease (CAVD) is the most common valvular heart disease in developed countries, yet no pharmacological therapy is available to slow or halt its progression. CAVD is driven by progressive calcification of aortic valve leaflets, in which myeloid cells play a central role. While macrophages have been implicated in CAVD pathogenesis, the contribution of their precursors, monocytes, remains poorly understood. We hypothesized that circulating monocytes acquire a pro-calcific and pro-inflammatory phenotype contributing to valve remodelling and CAVD progression.

**Methods:** We profiled circulating CD14^+^ monocytes from healthy volunteers (Vol), patients with CAVD, and without CAVD (NCAVD). Peripheral blood mononuclear cells (PBMCs) were isolated, and monocyte subpopulations were phenotyped by flow cytometry. Transcriptome profiling by RNA sequencing identified disease-associated gene signatures, which were validated by RT-qPCR. The CD14^+^ monocyte secretome was analysed using multiplex assays. Functional ability of CAVD-derived CD14^+^ monocytes to induce myofibroblastic transdifferentiation (MT) and osteoblastic differentiation (OD) of human valvular interstitial cells (VICS) was evaluated by immunocytochemistry and quantitative o-cresolphthalein complexone assays.

**Results:** In PBMCs, CAVD monocytes displayed a subpopulation shift, with an increased proportion of CD14⁺⁺CD16⁻ classical monocytes and a reduced CD14⁺CD16⁺⁺ non-classical monocyte levels. In CD14^+^ monocytes, transcriptomic analysis revealed upregulation of inflammation-related (*PDK4*) and calcification-related (*ATP2B1*) genes, alongside downregulation of immunomodulatory genes (*DDR1*, *IKBKE*). Secretome analysis showed reduced production of immunomodulatory and anti-osteoblastogenic cytokines (IL-4, CCL3) while promoting gene expression of factors promoting MT and OD in VICS. These alterations were associated with a marked monocyte-induced increase in αSMA and *OPN* expression in VICS and a two-fold increase in calcification.

**Conclusion:** We demonstrate for the first time that circulating monocytes from patients with CAVD exhibit enhanced pro-inflammatory and pro-calcific properties that may contribute to CAVD progression. Additionally, we identify dysregulated gene sets within these monocytes that represent potential novel therapeutic targets for CAVD.

## 4. Introduction

Calcific Aortic Valve Disease (CAVD) is the most common valvular heart disease worldwide ^1^. Its prevalence increases with age, affecting approximately 2% of individuals over 65 years and up to 10% of people aged over 80 years^2,3^. As an age-related condition, the growing proportion of elderly individuals in the population has made CAVD a major public health and economic challenge ^4,5^. CAVD is a severe disease, with a high mortality rate of approximately 70% following symptom onset in the absence of treatment ^6^. Currently, there is no effective pharmacological treatment available to prevent or halt the progression of CAVD. However, patients in advanced stages of the disease may undergo surgical aortic valve replacement (SAVR) or transcatheter aortic valve replacement (TAVR) ^7^. The pathogenesis of CAVD is initiated by endothelial injury and dysfunction, which are associated with disturbed blood flow ^8,9^. Subsequently, dysfunctional valvular endothelial cells (VECs) overexpress adhesion molecules, such as Intercellular adhesion molecule (ICAM-1) and vascular cell adhesion protein 1 (VCAM-1), facilitating monocyte adhesion and secrete inflammatory signals enhancing monocyte chemotaxis and infiltration into the aortic valve leaflets. Enhanced monocyte infiltration contributes to a pro-inflammatory microenvironment supporting CAVD development. Monocytes have thus emerged as both predictive biomarkers for disease advancement and promising therapeutic targets to prevent further valve degeneration^10^. Within this pro-inflammatory microenvironment, infiltrating monocytes polarize and differentiate into pro-inflammatory macrophages, thereby promoting valve inflammation. These macrophages secrete high levels of pro-inflammatory mediators, including interleukin (IL)-1β, IL-6, and tumor necrosis factor (TNF)-α, which promote the osteogenic differentiation of valvular interstitial cells (VICs) ^11,12^. Valvular inflammation is reported as a key regulator of CAVD development ^13,14^. Previous studies have demonstrated that chronic inflammatory infiltration by monocytes-macrophages, lymphocytes and mast cells, in aortic valve tissues promotes fibrosis and calcification^15,16^. Moreover, pro-inflammatory macrophages derived from infiltrating monocytes accumulate in the aortic valve leaflets and contribute to VIC fibrosis and calcification ^17,18^. In CAVD, VICs, the predominant cell type in aortic valve leaflets, play a central role in many local processes associated with valve calcification ^19,20^. Indeed, studies have shown that VICs undergo a phenotypic shift from a myofibroblast-like to an osteoblast-like morphology, a deleterious transformation that contributes to the development of calcifications in CAVD ^21^. Monocytes may be functionally divided into three main subpopulations: Cluster differentiation (CD)14^++^CD16^-^ classical monocytes (Mo1), CD14^++^CD16^+^ intermediate monocytes (Mo2), and CD14^+^ CD16^++^ non-classical monocytes (Mo3) ^22,23^. Emerging studies have demonstrated that elevated levels of circulating monocytes, principally CD14^++^ monocytes, are associated with severe CAVD ^24–26^. Therefore, we hypothesized that CAVD monocytes may be characterized by increased pro-inflammatory and pro-calcific capacities leading to increased local VICs activation, accelerating valve fibrosis and calcification and thus CAVD development. The objective of this study is to characterize the pro-inflammatory and pro-calcific properties of circulating monocytes from CAVD patients and to investigate their direct effects on VICs to better understand the cellular mechanisms driving valvular inflammation, fibrosis, and calcification during CAVD progression.

## 5. Methods

### Data Availability

The datasets generated and analysed during the current study are available from the corresponding author upon reasonable request.

## 6. Results

### Enriched CD14⁺⁺CD16⁻ monocyte subset in CAVD PBMCs is associated with pro-inflammatory profile

In our cohort, demographic variables and overall monocyte counts (Table 1) did not differ significantly between NCAVD and CAVD groups. However, flow cytometry revealed differences in monocyte subpopulations between NCAVD and CAVD group (clinical characteristics summarized in Table 2), on one hand, and Vol group on the other (Figure S1), consistent with previous descriptions ^24–26^. We observed an increase of CD3^+^CD20^-^ T lymphocytes in NCAVD and CAVD compared to Vol, however no differences were found in T cells or in CD3^-^CD20^+^ B cells between the three groups (Figure S2). NCAVD patients exhibited a higher percentage of CD14⁺⁺CD16⁻ Mo1 and a lower percentage of CD14⁺CD16⁺⁺ Mo3 compared to Vol. Additionally, the proportions of CD14⁺⁺CD16⁻CD86⁺ activated monocytes were increased, while CD14⁺CD16⁺⁺CD206⁺ monocytes were decreased in NCAVD patients. No significant differences were observed for CD14⁺⁺CD16⁺ Mo2 across groups. CAVD patients showed an even more pronounced increase than NCAVD patients in CD14⁺⁺CD16⁻ Mo1 and decrease in CD14⁺CD16⁺⁺ Mo3 compared to Vol. Moreover, the proportion of CD14⁺⁺CD16⁻CD86⁺ monocytes was further elevated in the CAVD group, while the reduction in CD14⁺CD16⁺⁺CD206^+^ monocytes mirrored that found in NCAVD patients (Figure 1). These findings suggest that monocytes in CAVD patients may be primed for an enhanced inflammatory response, thereby contributing to the pro-inflammatory environment and disease progression. While this inflammatory profile is not exclusive to CAVD patients, their monocytes tend to be more pro-inflammatory than those from the NCAVD group.

**Figure 1.**
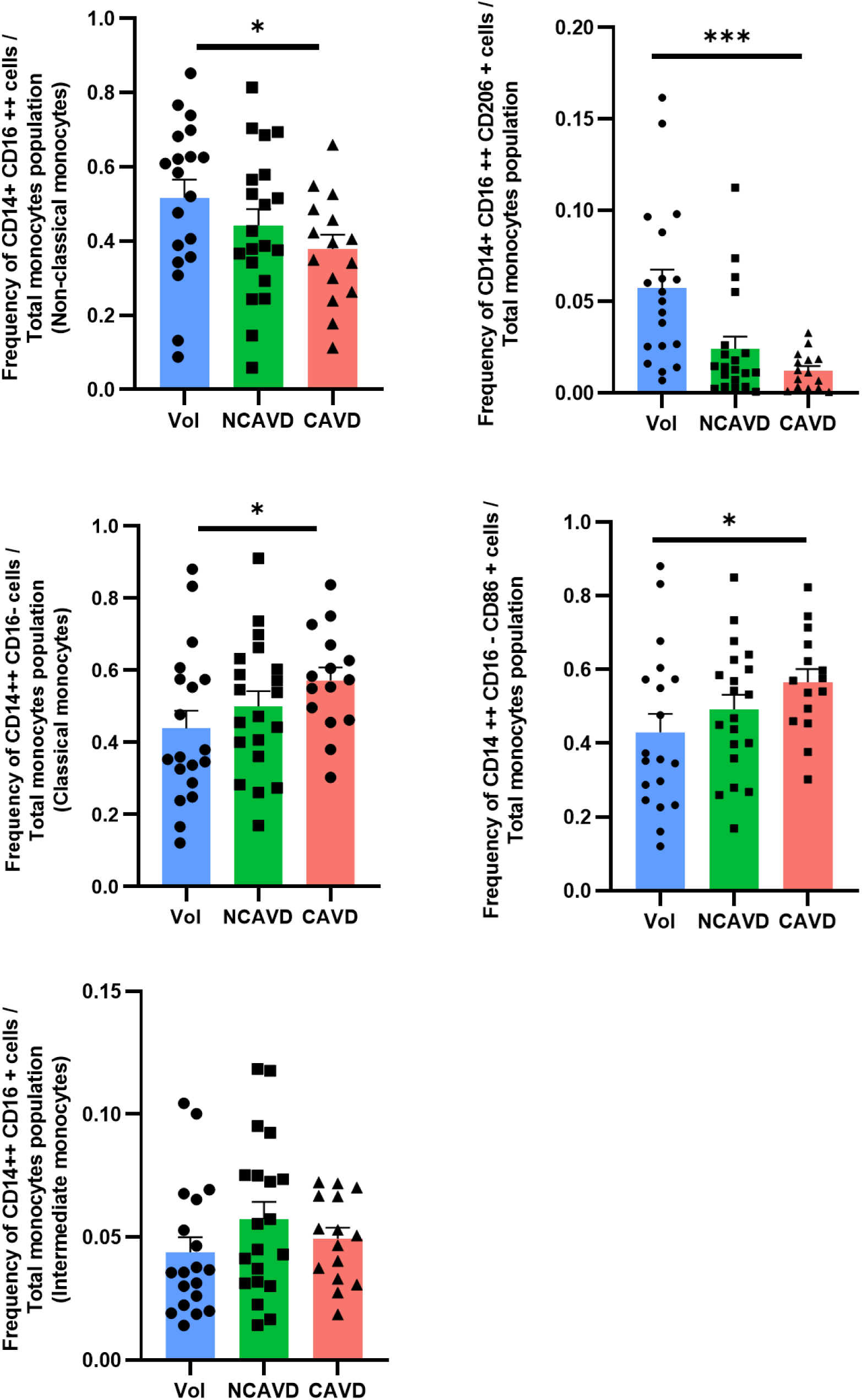
CD14⁺⁺CD16⁻ Mo1 enrichment in CAVD patients drives heightened inflammatory responsiveness. Flow cytometry analysis of peripheral blood monocytes from healthy volunteers (Vol, n=19), NCAVD (n=20), and CAVD patients (n=15). Classical (CD14⁺⁺CD16⁻), intermediate (CD14⁺⁺CD16⁺), and non-classical (CD14⁺CD16⁺⁺) monocytes subsets were identified, and their M1-like (CD86⁺) or M2-like (CD163⁺, CD206⁺) phenotypes assessed. Frequencies were expressed as % of total monocytes. Data were analysed by one-way ANOVA with Sidak’s multiple comparisons test (* p < 0.05; *** p < 0.001).

**Table 1.**
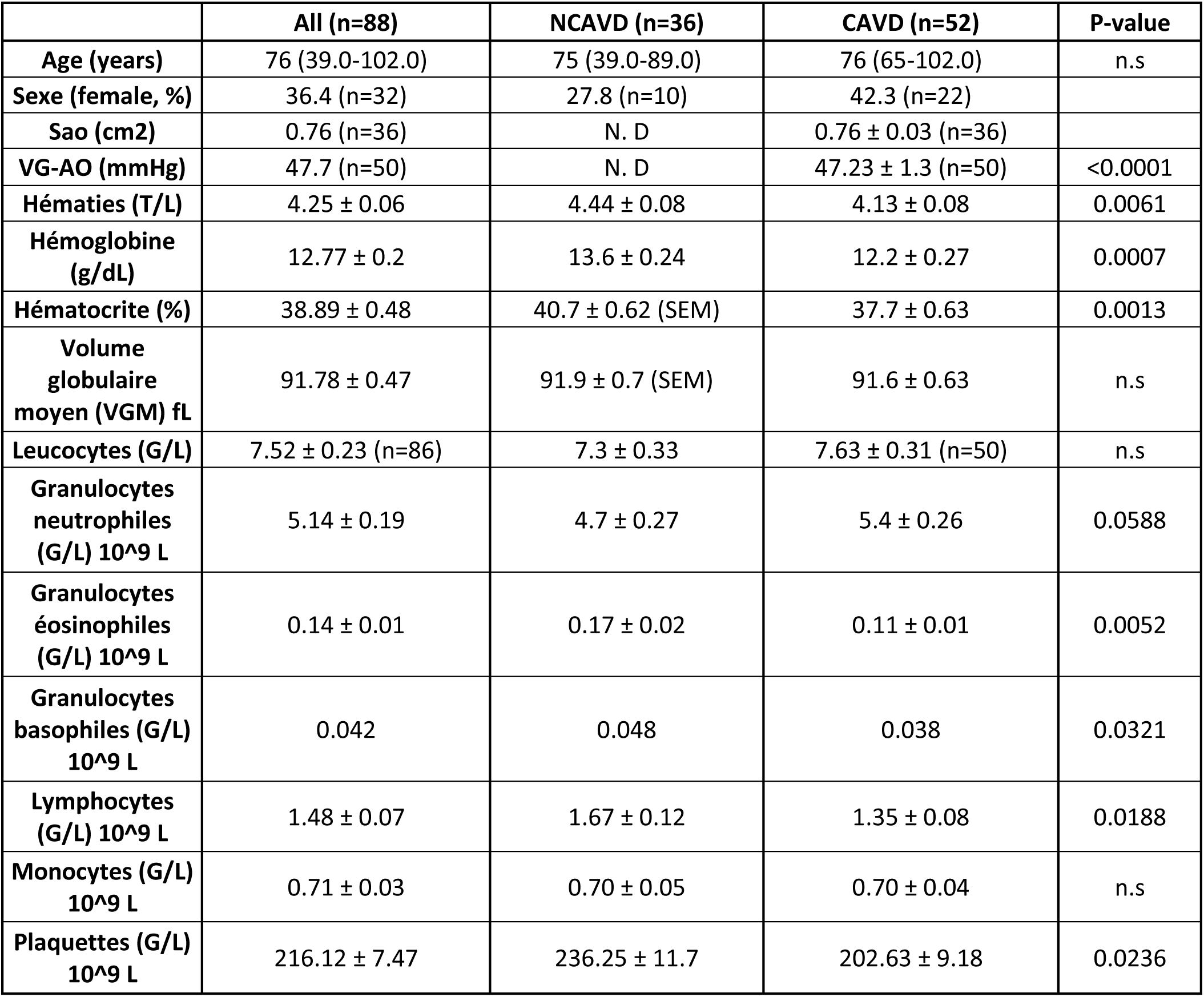
Clinical characteristics of patients. Data were collected from n = 36 out of 58 coronary artery disease patients without calcific aortic valve disease (NCAVD) and n = 52 out of 60 patients with severe calcific aortic valve disease (CAVD). Legend: N. D = No data; SAo = aortic valve area; LV–AO = transvalvular gradient measured as the pressure difference between the left ventricle (LV) and the aorta (AO). Statistical significance between NCAVD and CAVD was assessed using either a parametric or a non-parametric t-test, depending on the distribution determined by the Shapiro–Wilk normality test. p-value is indicated when significant, with ns for p > 0.05.

|  | All (n=88) | NCAVD (n=36) | CAVD (n=52) | P-value |
| --- | --- | --- | --- | --- |
| Age (years) | 76 (39.0-102.0) | 75 (39.0-89.0) | 76 (65-102.0) | n.s |
| Sexe (female, %) | 36.4 (n=32) | 27.8 (n=10) | 42.3 (n=22) |  |
| Sao (cm2) | 0.76 (n=36) | N. D | 0.76 ± 0.03 (n=36) |  |
| VG-AO (mmHg) | 47.7 (n=50) | N. D | 47.23 ± 1.3 (n=50) | <0.0001 |
| Hématies (T/L) | 4.25 ± 0.06 | 4.44 ± 0.08 | 4.13 ± 0.08 | 0.0061 |
| Hémoglobine (g/dL) | 12.77 ± 0.2 | 13.6 ± 0.24 | 12.2 ± 0.27 | 0.0007 |
| Hématocrite (%) | 38.89 ± 0.48 | 40.7 ± 0.62 (SEM) | 37.7 ± 0.63 | 0.0013 |
| Volume globulaire moyen (VGM) fL | 91.78 ± 0.47 | 91.9 ± 0.7 (SEM) | 91.6 ± 0.63 | n.s |
| Leucocytes (G/L) | 7.52 ± 0.23 (n=86) | 7.3 ± 0.33 | 7.63 ± 0.31 (n=50) | n.s |
| Granulocytes neutrophiles (G/L) 10<sup>9</sup> L | 5.14 ± 0.19 | 4.7 ± 0.27 | 5.4 ± 0.26 | 0.0588 |
| Granulocytes éosinophiles (G/L) 10<sup>9</sup> L | 0.14 ± 0.01 | 0.17 ± 0.02 | 0.11 ± 0.01 | 0.0052 |
| Granulocytes basophiles (G/L) 10<sup>9</sup> L | 0.042 | 0.048 | 0.038 | 0.0321 |
| Lymphocytes (G/L) 10<sup>9</sup> L | 1.48 ± 0.07 | 1.67 ± 0.12 | 1.35 ± 0.08 | 0.0188 |
| Monocytes (G/L) 10<sup>9</sup> L | 0.71 ± 0.03 | 0.70 ± 0.05 | 0.70 ± 0.04 | n.s |
| Plaquettes (G/L) 10<sup>9</sup> L | 216.12 ± 7.47 | 236.25 ± 11.7 | 202.63 ± 9.18 | 0.0236 |

**Table 2.** Clinical characteristics of patients with blood sample used for flow cytometry analysis. Data were collected from n = 20 out of 58 coronary artery disease patients without calcific aortic valve disease (NCAVD) and n = 16 out of 60 patients with severe calcific aortic valve disease (CAVD). Legend: N. D = No data; SAo = aortic valve area; LV–AO = transvalvular gradient measured as the pressure difference between the left ventricle (LV) and the aorta (AO). Statistical significance between NCAVD and CAVD was assessed using either a parametric or a non-parametric t-test, depending on the distribution determined by the Shapiro–Wilk normality test. p-value is indicated when significant, with n.s for p > 0.05.

|  | All (n=36) | NCAVD (n=20) | CAVD (n=16) | P-Value |
| --- | --- | --- | --- | --- |
| **Age (years)** | 73 (40.0-89.0) | 67 (40.0-87.0) | 79 (46.0 - 89.0) | 0.0018 |
| **Sexe (female, %)** | 46.9% (n=15) | 35.0% (n=7) | 50.0% (n=8) |  |
| **Sao (cm2)** | 0.86 (n=7) | N. D | 0.86 (n=7) | <0.0001 |
| **VG-AO (mmHg)** | 49.3 (n=15) | N. D | 49.3 (n=15) | <0.0001 |

### CAVD monocytes display a prominent pro-inflammatory gene expression profile

Whole-transcriptome RNA sequencing of freshly isolated monocytes was performed to investigate the molecular mechanisms driving the enrichment of CD14⁺monocytes in CAVD patients. The analysis revealed 548 upregulated and 391 downregulated genes in CAVD monocytes relative to NCAVD CD14^+^ monocytes (Figure 2A). We performed Gene Ontology (GO) analysis to identify biological processes enriched among the differentially expressed genes in monocytes from CAVD patients. Compared with CD14^+^ monocytes from Vol, CD14^+^ monocytes from both NCAVD and CAVD patients exhibited upregulation of genes such as C-X3-C motif chemokine receptor 1 (*CX3CR1*) and Toll-like receptor 2 (*TLR2*), which are associated with enhanced inflammation and innate immune responses (Figure S3D and F). Conversely, genes related to immunomodulatory functions, including Krüppel like factor 4 (*KLF4*) and Cytokine like 1 (*CYTL1*), were downregulated in these patient groups (Figure S3E and F). Particularly for CAVD CD14^+^ monocytes, upregulated genes, such as Pyruvate dehydrogenase kinase 4 (*PDK4*), ATPase plasma membrane Ca2^+^ transporting 1 (*ATP2B1*), were enriched in GO terms related to enhanced calcification and migration properties (Figure 2B, D and F). Conversely, downregulated genes, including Inhibitor of nuclear factor kappa-B kinase subunit epsilon (*IKBKE*), Discoidin domain receptor tyrosine kinase 1 (*DRR1*), were associated with immunomodulatory GO biological processes (Figure 2C, E and F). These findings suggest that monocytes from CAVD patients exhibit an enhanced pro-inflammatory profile, consistent with the observed increase in the CD14⁺⁺CD16^-^ Mo1 subset. This dysregulated transcriptomic signature may underlie augmented calcification and chemotaxis processes, alongside impaired immunomodulatory functions of circulating monocytes in CAVD patients.

**Figure 2.**
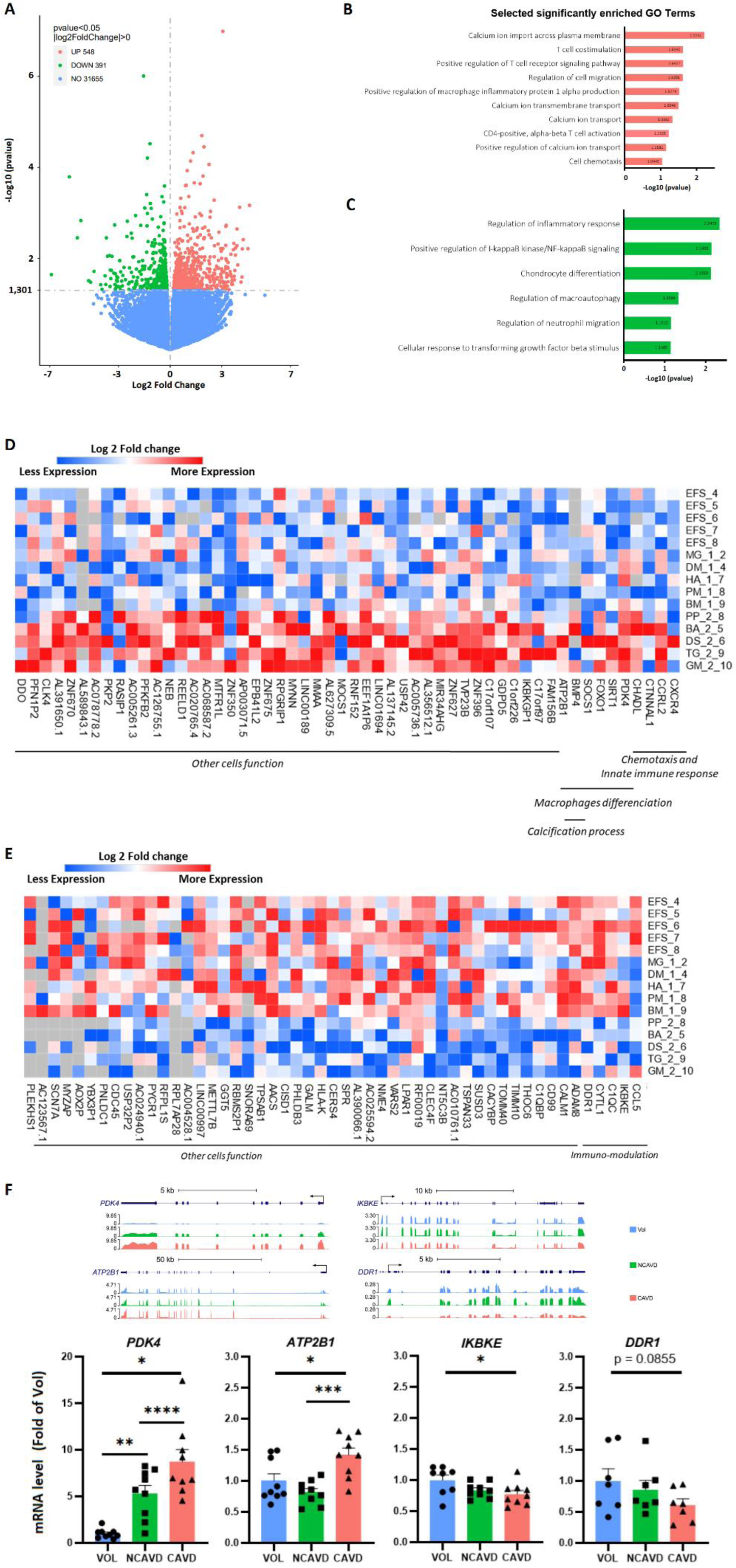
Transcriptomic profiling reveals dysregulated genes signature in CAVD CD14^+^monocytes. A) Volcano plot showing upregulated (red) and downregulated (green) genes from combined data of five donors in the NCAVD group and five donors in the CAVD group. Genes were considered statistically significant at adjusted p-value < 0.05, calculated using a negative binomial model (Novogene). (B, C) Bar graphs representing Gene Ontology (GO) Biological Processes enriched among upregulated (B) and downregulated (C) genes. (D, E) Heatmaps of selected significantly upregulated (D) and downregulated (E) genes identified by RNA-seq. (F) mRNA expression of genes of interest was measured by RT-qPCR and normalized to B2M. Sample sizes were as follows: for *ATP2B1* and *PDK4*, n = 9 per group; for *IKBKE*, n = 9 in the NCAVD and CAVD groups and n = 7 for healthy volunteers (Vol); for *DDR1*, n = 7 per group. Statistical analysis was performed using one-way ANOVA with Sidak’s post-hoc test; outliers were removed. P-values are indicated as: *p < 0.05, **p < 0.01, ***p < 0.001, ****p < 0.0001.

### Impaired immunomodulatory cytokine secretion by CAVD monocytes reflects an enhanced pro-inflammatory phenotype

As myeloid cells, including monocytes, may communicate with VICs through chemokine and cytokine secretion to induce myofibroblastic transdifferenciation (MT) and osteogenic differentiation (OD), we focused on their transcriptomic expression. Compared with CD14^+^ monocytes from healthy volunteers, NCAVD CD14^+^ monocytes displayed reduced expression of C-C motif ligand (*CCL*)3 and *IL4*. A similar reduction was also observed in CAVD monocytes (Figure 3A). These results were further validated by high-throughput Luminex secretome profiling, which demonstrated diminished secretion of both cytokines in NCAVD CD14^+^ monocytes, and an even greater reduction in CAVD CD14^+^ monocytes (Figure 3B). We found a decreased secretion of CCL21 by NCAVD CD14^+^ monocytes compared to Vol monocytes with an even greater reduction found in CAVD monocytes. Notably, in CAVD CD14^+^ monocytes, we also found diminished secretion of C-X-C motif ligand (CXCL)9, CCL17 and CCL22 chemokines that play key roles in mediating communication with the adaptive immune system through lymphocyte interactions (Figure S4). These findings demonstrate that monocytes from CAVD patients contribute to a pro-inflammatory microenvironment through decreased immunomodulatory cytokine secretion, which may amplify inflammatory responses and promote disease development.

**Figure 3.**
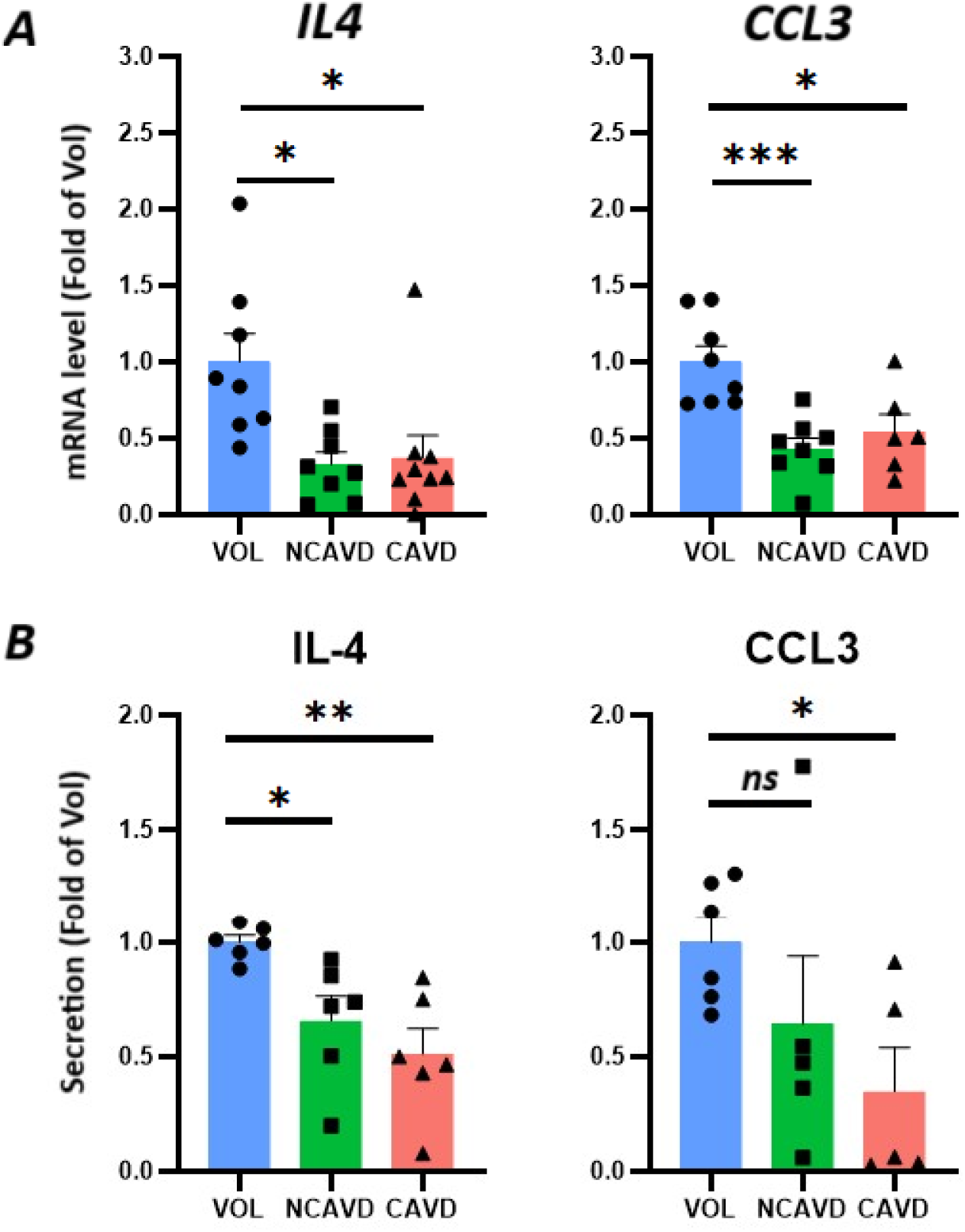
CAVD CD14^+^monocytes secretome is depleted in immunoregulatory and anti-osteogenic cytokines. *IL4* and *CCL3* mRNA expression were quantified by RT-qPCR (A) and normalized to the housekeeping gene *B2M*. Sample sizes were as follows: for *IL4*, n = 8 per group; for *CCL3*, n = 8 in the NCAVD and healthy volunteers (Vol) groups and n = 6 for CAVD group. (B) IL-4 and CCL3 CD14^+^ monocytes secretion were quantified by multiplex immunoassays. Sample sizes were as follows: for IL-4, n = 6 per group; for CCL3, n = 6 in the healthy volunteers (Vol) group and n = 5 for NCAVD and CAVD groups. Asterisks (*) indicate statistically significant differences compared to the healthy volunteer condition (blue) as determined by One-Way ANOVA followed by Sidak’s post-hoc test. Outliers were removed. p-values are indicated as follows: *p < 0.05, **p < 0.01, ***p < 0.001.

### Pro-inflammatory secretome of CAVD monocytes drives VIC myofibroblastic transition and osteogenic differentiation

To investigate the functional impact of CAVD CD14^+^ monocyte secretome on VIC MT and OD, we exposed VICs to conditioned media from CAVD CD14^+^ monocytes cultured for 3 days. VICs treated with CAVD CD14^+^ monocyte-conditioned media exhibited a more fibrotic phenotype, with increased number of Alpha smooth muscle actin (αSMA) ^+^cells compared to VICS exposed to non-conditioned media (Figure 4A and B). This observation was confirmed at the mRNA level, including an increase in fibroblastic gene expression such as Tenascin C (*TNC*) and Tissue inhibitor of metalloproteinases (*TIMP1*) (Figure 4C). Building on our observation that conditioned media from CAVD monocytes promote a pro-fibrotic phenotype in VICSs, we next examined whether these secretions also drive osteogenic differentiation and calcium deposition. We observed that treatment with CAVD CD14^+^ monocyte-conditioned media significantly increased the number of Osteopontin (OPN)^+^ cells and calcium deposits compared to non-conditioned media (Figure 5A and B). This observation was confirmed by increased expression of osteogenic genes, Alkaline phosphatase (*ALPL*) and Osteoprotegerin (*OPG*) (Figure 5D). In agreement, CAVD CD14^+^ monocyte-conditioned media exposed VICs produced more calcium deposits (Figure 5E). Interestingly, we found in VICs treated with Vol CD14^+^ a marked upregulation of *TIMP1* along with a trend toward increased *TNC* gene expression (Fig. S5A), suggesting that secretions from Vol CD14⁺ monocytes exert a modest effect on VIC myofibroblastic transdifferentiation. Regarding osteogenic differentiation and calcium deposition, treatment with Vol CD14⁺ monocyte secretions had no significant effect on calcium deposits (Fig S5 B) compared with non-conditioned medium condition. No significant increase in *OPG* gene expression was detected. In contrast, *ALPL* expression was significantly upregulated, which may be attributable to the use of unhealthy VICs in our experiment (Fig S5A). Altogether, these findings indicate that the monocyte secretome, especially from CAVD monocytes, may promote VIC myofibroblast transition and subsequent osteogenic differentiation during CAVD progression

**Figure 4.**
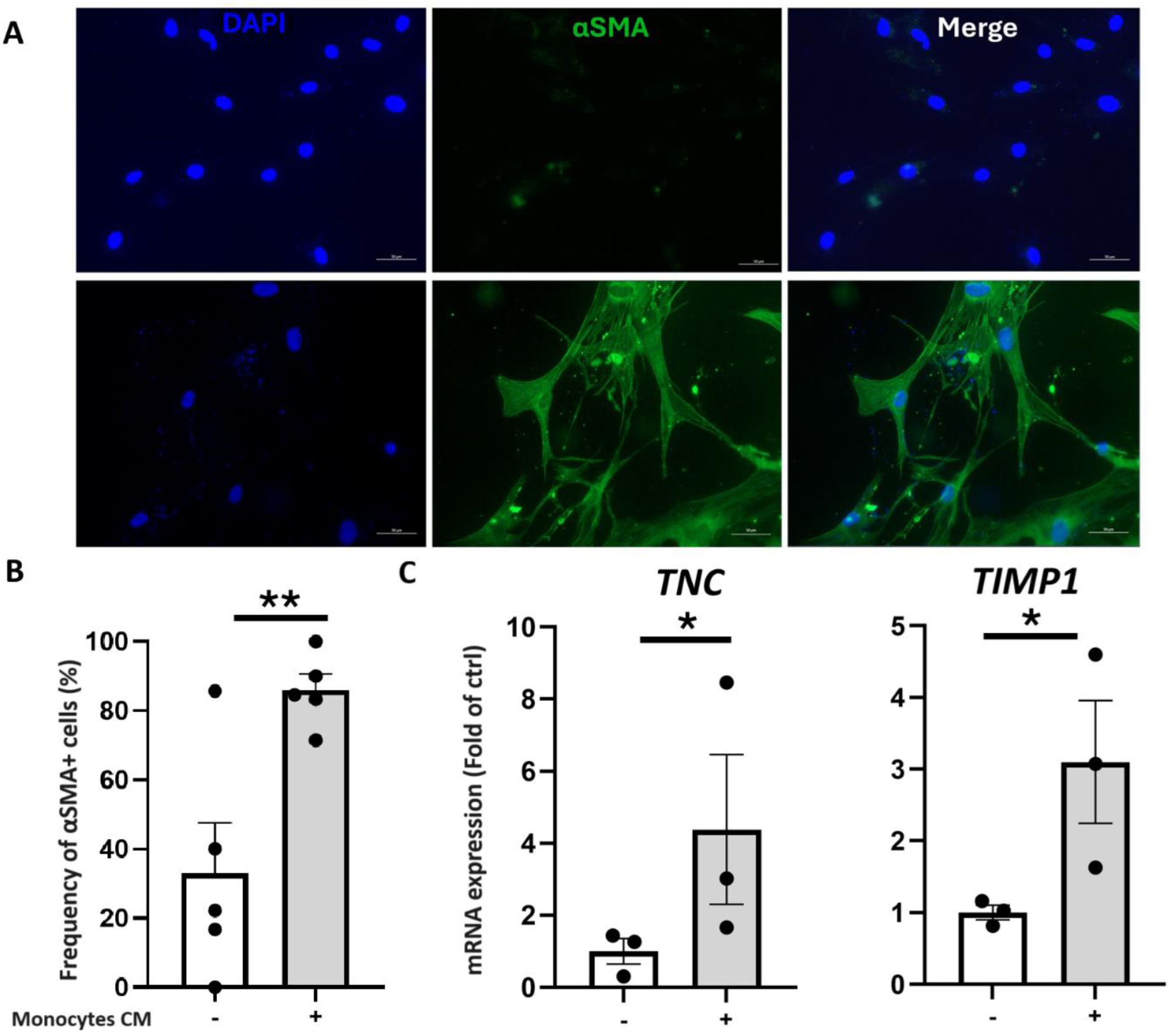
CAVD CD14^+^ monocytes promote myofibroblastic transition and pro-fibrotic gene expression in VICS. Representative pictures of α-SMA positive VICS exposed to CAVD monocytes conditioned media or non-conditioned media for 3 days (A). Nuclei were counterstained with DAPI (blue) and the fibrosis marker α-SMA is shown in green. Scale bars: 50 μm. Quantification of α-SMA–positive VICS (B). Five images per condition were acquired using a Zeiss Axio Imager Z1 fluorescence microscope and analyzed with Zeiss Zen software. *TNC* and *TIMP1* gene expression in VICS (C) measured by RT-qPCR (n = 3 per condition) after treatment with CAVD monocytes conditioned media or non-conditioned media for 3 days. Gene expression is normalized to the housekeeping gene *B2M*. Statistical analysis was performed using Student’s t-test. p-values are indicated as follows: *p < 0.05, **p < 0.01.

**Figure 5.**
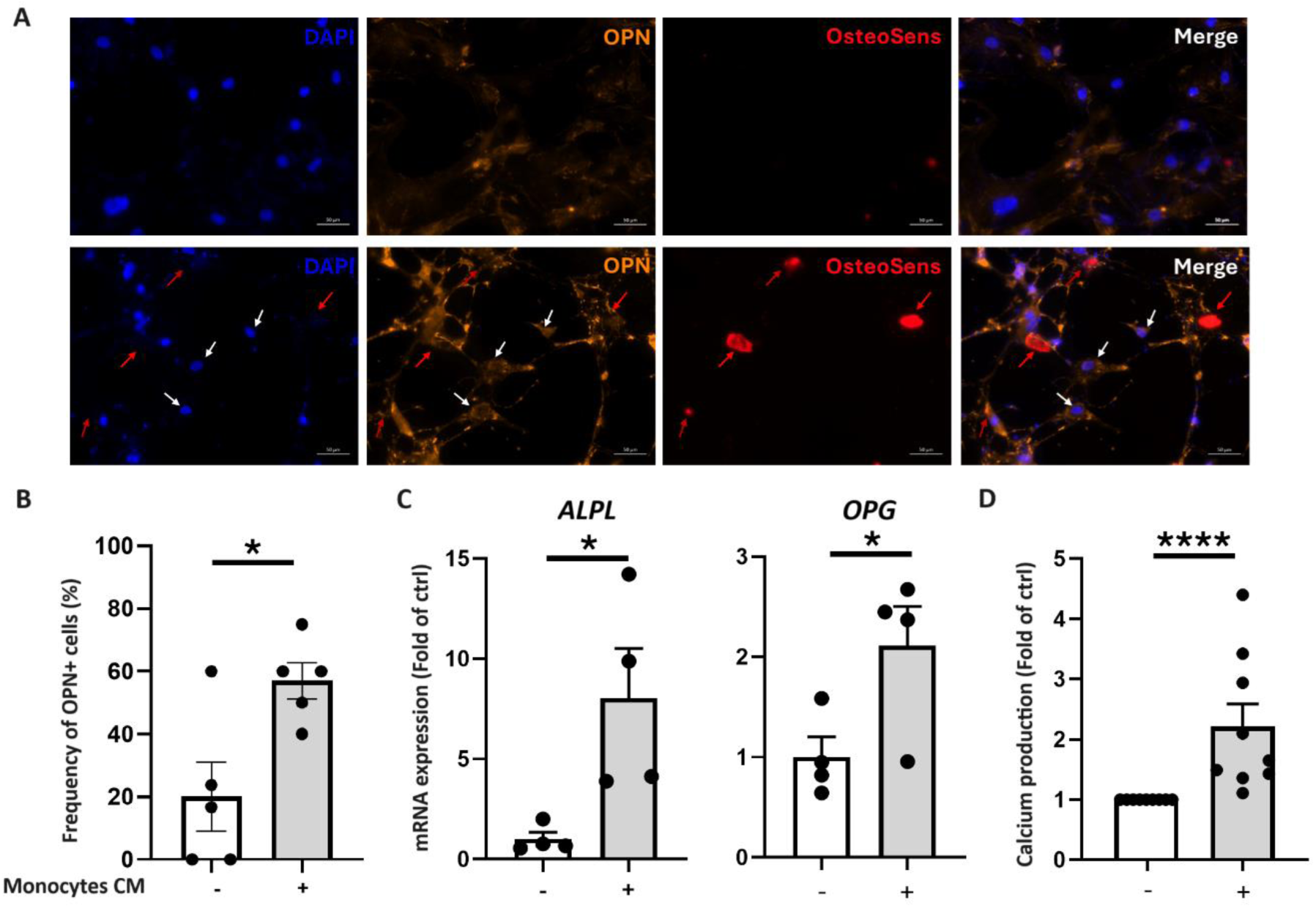
Conditioned media from CAVD CD14^+^ monocytes promote VICS osteogenic transition and calcium production. Representative pictures of osteopontin (OPN) positive VICS and calcium production (OsteoSens^TM^) after exposition to CAVD monocytes conditioned media or non-conditioned media for 7 days (A). Nuclei are stained with DAPI (blue) and calcification markers OPN and OsteoSens are shown in orange and red respectively. Scale bars: 50 μm. Quantification of OPN–positive VICS (B). Five images per condition were acquired using a Zeiss Axio Imager Z1 fluorescence microscope and analysed with Zeiss Zen software. *ALPL* and *OPG* gene expression in VICS (C) measured by RT-qPCR (n = 4 per condition) after treatment with CAVD monocytes conditioned media or non-conditioned media for 7 days. Gene expression is normalized to the housekeeping gene *B2M*. Hydroxyapatite crystals(D) were quantified using O-cresolphtalein assay (n=9 for each condition). Statistical analysis was assessed with Mann-Whitney test. The p values are depicted as follows: * p<0.05 and **** p<0.0001. White arrows show OPN+ cells and red arrows calcifications.

## 7. Discussion

In CAVD, endothelial injury of aortic valve leaflets represents the initial trigger for immune cell infiltration. Circulating monocytes are among the first myeloid cells to be recruited to the damaged valve leaflet, where they adhere and differentiate into pro-inflammatory classical macrophages. These macrophages secrete cytokines including IL-1β, IL-6, and TNF-α, thereby amplifying the local inflammatory responses ^27^. These macrophages mainly arise from CD14^++^CD16^-^ Mo1 and CD14^++^CD16^+^ iM02^28^. Furthermore, patients with severe CAVD have been shown to exhibit higher levels of circulating CD14^+^ monocytes compared to healthy individuals ^24^. Our findings highlight that in the peripheral blood of CAVD patients, the proportion of CD14^++^CD16^-^ Mo1 is higher than CD14^+^CD16^++^ Mo3 compared to Vol. There is no change observed in the Mo2 subset. Moreover, we found higher proportion of CD14^++^CD16^-^CD86^+^ and reduced levels of CD14^+^CD16^++^CD206^+^ monocytes in CAVD patients. Whereas Mo1 participate in the pro-inflammatory responses, Mo3 have a role in tissue repair and resolution of inflammation. Mo2 may be involved in both pro-inflammatory responses and tissue repair ^29,30^. Furthermore, CD86 and CD206 markers are associated with an activated pro inflammatory M1-like state and immunomodulatory M2-like macrophages phenotypes, respectively ^31^. Thus, higher proportion of Mo1 associated with reduced proportion of Mo3 and unchanged proportion of Mo2 suggests that imbalanced systemic monocyte polarization in peripheral blood of CAVD patients contributes to sustain pro-inflammatory environment allowing continuous progression of CAVD and calcification.

Aside from TAVR and SAVR, no effective pharmacological treatments currently exist to slow or reverse CAVD. Identifying new therapeutic targets is therefore crucial to developing novel pharmacological strategies. Our CAVD CD14^+^ monocyte profiling, at both transcriptomic and functional levels, revealed elevated pro-inflammatory capacity as compared to in healthy volunteers. Among the differentially expressed genes we identified several associated with chronic inflammatory diseases including CAVD (Figure supp S3 A, B, C, D, E and F). To find specific CAVD-specific dysregulated genes we compared NCAVD and CAVD CD14^+^ monocytes transcriptomes. We found that genes differentially expressed in CAVD were associated with pro-inflammatory and immunomodulatory biological processes. Among these genes, we identified *PDK4* and *ATP2B1*. PDK4 is involved in regulating cellular metabolism by inhibiting the pyruvate dehydrogenase complex (PDC) ^32,33^. Dysregulation of the PDC can have important effects on cellular metabolism exacerbating the progression of diabetes, cardiovascular diseases, and the development of cancers ^34–36^. Increased expression of *PDK4* in CAVD monocytes may indicate a metabolic adaptation consisting of a shift from oxidative phosphorylation to glycolysis, frequently observed in inflammatory settings ^37,38^. *ATP2B1* encodes a calcium pump allowing maintenance of intracellular calcium homeostasis ^39,40^. Upregulation of *ATP2B1* in CAVD monocytes implies alteration in calcium handling that may contribute to their pro-inflammatory phenotype and directly impact the calcification process by influencing the deposition and regulation of calcium ions within the valvular tissue observed in CAVD.

Furthermore, we identify *IKBKE* and *DRR1*. IKBKE is a protein kinase involved in the regulation of the nuclear factor-kappa B (NF-κB) signalling pathway, which plays a critical role in inflammation, immunity, and cell survival ^41,42^. This gene is known to regulate innate immune response, and its dysregulation has been associated with autoimmune disorders, inflammatory diseases, and cancer ^43–45^. Decreased expression of *IKBKE* in CAVD monocytes indicates a potential impairment in immunomodulatory processes. *DDR1* encodes a receptor tyrosine kinase that binds to collagen and plays a role in cell adhesion, migration, and tissue remodeling ^46,47^. Several studies have demonstrated that dysregulation of DDR1 contributes to fibrotic diseases, cancer metastasis, and inflammatory responses ^48–50^. In CAVD, decreased expression of *DDR1* in monocytes suggests a potential disruption in extracellular matrix remodeling and tissue repair processes, which are crucial for maintaining valve homeostasis and preventing calcification.

In addition, we found that CAVD CD14^+^ monocytes exhibit altered secretion patterns. We observed decreased secretion of CCL3, an anti-osteoblastogenic cytokine, and IL-4, an immunomodulatory cytokine, in agreement with reduced gene expression levels. CCL3 Or macrophage inflammatory protein-1 alpha (MIP-1α) regulates the recruitment and activation of immune cells such as monocytes, macrophages, and T cells ^51^. Moreover, CCL3 promotes recruitment and activation of osteoclasts, the cells responsible for bone resorption, which affects osteoblast activity and bone remodeling ^52–54^. IL-4 is an anti-inflammatory cytokine that plays a central role in regulating immune responses. It is produced by activated T cells, mast cells, and basophils. IL-4 promotes the differentiation of naive T cells into Th2 cells, enhances B cell proliferation and induce macrophages polarization towards an anti-inflammatory phenotype ^55–57^. Additionally, our analysis revealed decreased secretion of CXCL9 and CCL21, cytokines involved in T cell recruitment ^58,59^, as well as reduced secretion of CCL17 and CCL22, cytokines associated with the Th2 response^60–62^ (Figure S4). Taken together, these results suggest altered capacity of CAVD monocytes in modulating immune cell recruitment and polarization within the valve microenvironment.

As monocytes communicate with other cell type through their secretome, we investigated whether their enriched pro-inflammatory secretions influenced VIC MT and OD. On one hand, we observed an upregulation of *TIMP1* and *TNC* gene expression in VICs treated with conditioned media from CAVD CD14^+^ monocytes. Accordingly, our immunocytochemistry results suggested that VICs had a pro fibrotic αSMA+ phenotype^63–65^. TIMP1 has a role in regulating extracellular matrix remodeling by inhibiting matrix metalloproteinase (MMP)s, thus promoting fibrosis ^66,67^. In CAVD, *TIMP1* upregulation can enhance fibrosis by preventing MMP-mediated extracellular matrix (ECM) degradation, leading to excessive ECM accumulation thus stiffening of the tissue. TNC is an extracellular matrix glycoprotein that contributes to tissue remodeling and fibrosis through interactions with cell surface receptors such as the epidermal growth factor receptor (EGF) and integrins in the ECM ^68–70^. In CAVD, increased expression of *TNC* maintains the pro-inflammatory response and tissue remodeling that contribute to valve fibrosis and calcification ^71^. On the other hand, we found increased expression of *ALPL* and *OPG*, genes related to calcification^72,73^ in VICs exposed to conditioned media from CAVD monocytes. In agreement, immunocytochemistry analysis revealed a greater number of OPN+ VICs and increased calcification deposits after stimulation with secretome from CAVD CD14^+^ monocytes. In CAVD, increased expression of *ALPL* in VICs can enhance mineralization of the valve tissue, contributing to the development of calcification. OPG is a decoy receptor for the receptor activator of NFkB ligand (RANKL) and regulates bone remodeling by inhibiting RANKL-mediated osteoclastogenesis and bone resorption ^74,75^. In CAVD, *OPG* overexpression can impact the balance between bone formation and resorption therefore calcification in the aortic valve leaflets ^76^. OPN is a multifunctional protein involved in bone metabolism and mineralization ^77,78^. In CAVD, upregulation of OPN is associated with the progression of valvular calcification and is considered a marker of disease severity ^79,80^.

The interaction between ALPL, OPG, and OPN in VICs might contribute to the dysregulated mineralization promoting calcification within the aortic valve. Overall, our findings suggest that CAVD monocytes influence VICs by promoting both fibrosis and calcification processes.

The main limitations of our study include the limited sample size, the use of unhealthy VICs and the complexity of *in vivo* cellular interactions that cannot be fully replicated *in vitro.* Further research is needed to elucidate the exact mechanisms through which monocytes influence VICs and to identify therapeutic targets to mitigate inflammation and calcification in CAVD.

## Conclusion

We demonstrated for the first time that circulating CAVD monocytes have a higher pro-inflammatory capacity and promote VICs fibrosis and calcification associated with aortic valve leaflets calcification and subsequent CAVD development. Moreover, we identified a set of dysregulated genes which may represent new therapeutic targets to prevent and/or limit disease development. Further studies are necessary to better understand how to regulate immune cell activity, including monocytes and lymphocytes, to reduce CAVD development.

## 7. Acknowledgments

The authors would like to thank the patients, families, and staff at the cardiology department in Rouen university hospital. They also thank Dr Virginie Tardif for her expertise advice during preparation of flow cytometry analyses.

## 9. Disclosure

The authors declare no conflict of interest.

### Human peripheral blood collection

In accordance with French legislation, all patients received an information sheet regarding the protocol and provided oral consent for the collection of an additional blood volume during their routine care, as well as for the processing of their personal data (age, sex, etc.). Peripheral blood was collected during coronary procedures for NCAVD patients, prior to scheduled interventions for CAVD patients, and during blood donation for healthy volunteers. Samples were drawn into four BD Vacutainer EDTA K2 tubes (Beckton Dickinson, Cat#367862). This study did not consider sex-based or race/ethnicity-based differences. Inclusion criteria for CAVD patients were: (1) age equal or greater than 18 years old, (2) admission to hospital for a coronary angiography (3) severe aortic stenosis (AVA < 1 cm2 and/or LV-AO > 40 mmHg) (4) affiliated to medical health insurance. Exclusion criteria were: (1) Infectious disease, (2) Obesity (BMI> 30kg/m2), (3) Haematological pathologies, (4) Previous TAVR, (5) Anaemia and (6) Inflammatory and autoimmune pathologies. Coronary patients had no aortic stenosis, and similar exclusion criteria as CAVD patients. In-hospital data were recorded in a dedicated database. The EFS (Établissement Français du Sang) is a public administrative body responsible for the collection, preparation, qualification, and distribution of labile blood products (blood, plasma, and platelets) for transfusion purposes in France. For its non-transfusion-related activities, EFS is authorized to collect, process, store, and provide blood or its components for educational or research purposes, excluding any therapeutic use. Only adults (aged 18 years and older) are eligible to donate blood. Blood collection is performed exclusively with the donor’s written consent. The physician informs the donor of the significance of biological samples for advancing medical research.

### VICS isolation and culture

Aortic valves were cut into small pieces and digested using collagenase I (Fisher Scientific, Cat#10114532). Isolated VICS were initially cultured in Smooth Muscle Cell Growth Medium (PromoCell, Cat#C-22162). Following the first passage, cells were maintained in Dulbecco’s Modified Eagle Medium (DMEM; Gibco, Cat#41966-029) supplemented with 10% fetal bovine serum (FBS; Gibco, Cat#10500-064) and 1% penicillin/streptomycin (Sigma-Aldrich, Cat#P4333). After one or two passages, VICSs were seeded in 6-well plates for RNA extraction and in 4-well plates for immunocytochemistry, and cultured for at least 5 days. Once confluent, VICSs were treated with patient-derived monocyte-conditioned media.

### Flow cytometry

Peripheral blood mononuclear cells (PBMCs) were isolated from the peripheral blood of NCAVD, CAVD, and healthy volunteers. Blood samples were collected in EDTA-containing tubes and processed using density gradient centrifugation over a Ficoll. Briefly, 500 µl blood was diluted 1:2 with sterile PBS and carefully layered over 1 mL of Ficoll in a 15 mL Falcon tube. Samples were centrifuged at 400 × g for 30 minutes at room temperature with no brake. Following centrifugation, PBMCs layer was collected, washed, and resuspended in FACS buffer (PBS supplemented with 2% FBS and 1 mM EDTA). To prevent non-specific antibody binding, a human Fc receptor blocking reagent was added to the cell suspension and incubated for 15 minutes at 4 °C. PBMCs were stained for 11-color flow cytometry analysis across 55 samples (NCAVD, n = 20; CAVD, n = 15; healthy volunteers, n = 20). Each staining panel included the following lineage markers: CD20, CD3, CD4, CD8, and CD25, allowing for the exclusion of B and T lymphocytes. Monocyte subsets were identified using CD14 and CD16. Additionally, CD86, CD163, and CD206 were used to assess monocyte polarization states—towards unpolarized, pro-inflammatory, or immunomodulatory macrophage phenotypes, respectively. Monoclonal antibodies (mAbs) were added to the cell suspension, and samples were incubated for 20 minutes on ice, protected from light. Optimal mAb concentrations were determined by prior titration experiments. Cells were then washed once with FACS buffer and fixed in 1% paraformaldehyde (PFA) for 15 minutes at 4 °C. After a final wash, cells were resuspended in FACS buffer and stored at 4 °C in the dark until flow cytometry analysis.

### RNA sequencing

Sequencing was performed using 150 nucleotide paired-end reads (PE150) on an Illumina NovaSeq 6000 platform. Reads were aligned to the human genome (hg38) and transcript assembly GRCh38 from Ensembl using HISAT2 version 2.0.5. Transcript quantification was conducted with featureCounts from the Subread package. Differential expression analysis, following normalization, was performed using the DESeq2 Bioconductor package version 1.22.1 in R version 3.5.1. Genes significantly up- or downregulated (adjusted p-value < 0.05) were submitted to DAVID Gene Ontology analysis version 6.7. Expression values (FPKM) for genes within selected GO categories were retrieved to generate heat maps. Heat maps were produced using Cluster version 3.0 and Java TreeView version 1.1.6r4.

### RT-qPCR

The following primers were used for RT-qPCR:

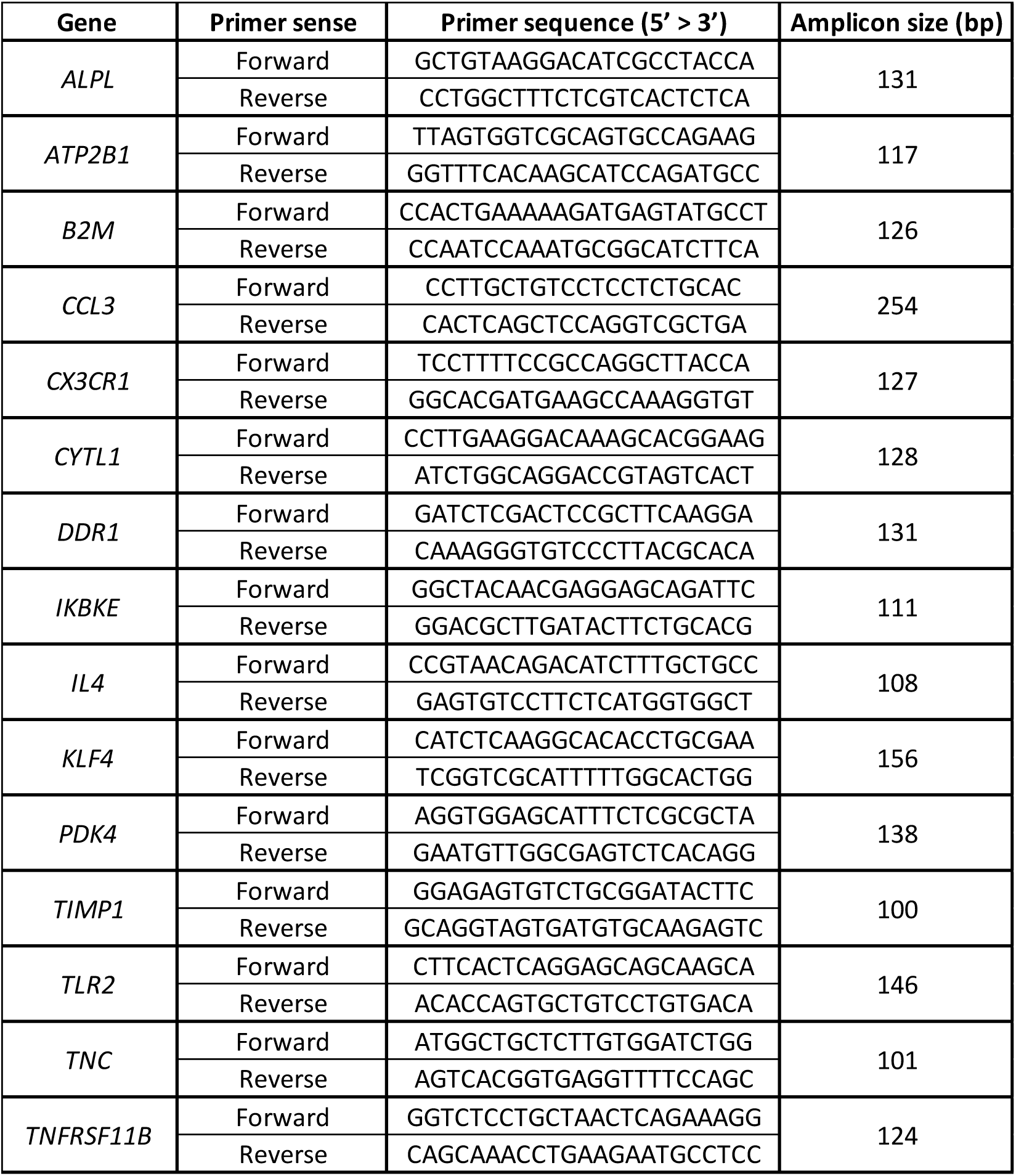

### O-cresolphthalein complexone assay

VICs were washed with 1X PBS and incubated overnight at 4 °C with 0.6 N HCl for decalcification. The resulting supernatant was collected and transferred to a 96-well plate. Calcium content was quantified by measuring absorbance at 565 nm using a Spark® Multimode Microplate Reader (Tecan). The calcium concentration was determined using a standard curve generated by linear regression.

### Antibodies and reagents

Flow cytometry:

| Antigen | Fluorochrome | Source and Catalog # |
| --- | --- | --- |
| CD20 | AF700 | Biolegend clone 2H7, Cat#302322 |
| CD3 | PerCP/Cy5.5 | Sony clone UCHT1, Cat#2102150 |
| CD4 | APC/Cy7 | Sony clone OKT4, Cat#218090 |
| CD8a | BV785 | Sony clone RPA-T8, Cat#2105230 |
| CD25 | FITC | Sony clone BC96, Cat#2113020 |
| CD14 | PE/Cy7 | eBioscience clone 61D3, Cat#25-0149-41 |
| CD16 | PE | eBioscience clone B73.1, Cat#12-0167642 |
| CD86 | BV605 | Sony clone IT2.2, Cat#2127145 |
| CD163 | APC | Life technologies clone GHI/61, Cat#A15723 |
| CD206 | eFluor450 | eBiosciences clone 19.2, Cat#48-2069-41 |
| Fixable Viability Dye (L/D) | UV | Invitrogen, Cat#L23105 |

Immunocytochemistry:

| Antigen | Supplier | Cat# | Species reactivity | Host | Dilution | [C] (µg/mL) |
| --- | --- | --- | --- | --- | --- | --- |
| Alpha SMA-FITC | Sigma Aldrich | F3777 | Human | Mouse | 1/200 | 14 |
| Osteopontin | Abcam | Ab8448 | Human | Rabbit | 1/500 | 164 |

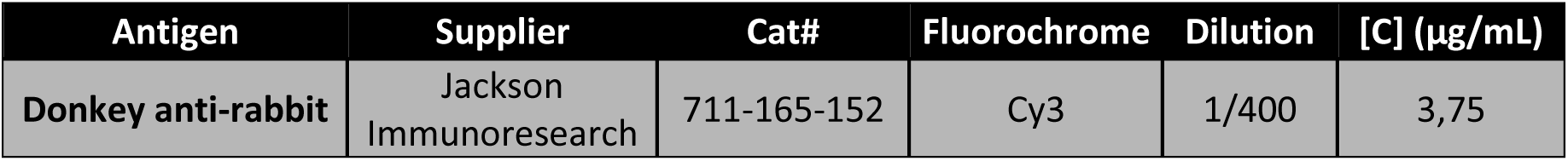

| Antigen | Supplier | Cat# | Fluorochrome | Dilution | [C] (µg/mL) |
| --- | --- | --- | --- | --- | --- |
| Donkey anti-rabbit | Jackson Immunoresearch | 711-165-152 | Cy3 | 1/400 | 3,75 |

## Abbreviation

αSMA: Alpha smooth muscle actin
ALPL: Alkaline phosphatase
ATP2B1: ATPase plasma membrane Ca2+ transporting 1
CAVD: Calcific aortic valve disease
CCL: C-C motif ligand
CD: Cluster differenciation
CXCL: C-X-C motif ligand
CX3CR1: C-X3-C motif chemokine receptor 1
CYTL1: Cytokine like 1
DDR1: Discoidin domain receptor tyrosine kinase 1
ECM: Extracellular matrix
EGF: Epidermal growth factor
GO: Gene Ontology
ICAM-1: Intercellular adhesion molecule 1
IKBKE: Inhibitor of nuclear factor kappa-B kinase subunit epsilon
IL: Interleukin
KLF4: Krüppel like factor 4
MIP-1α: Macrophage inflammatory protein-1 alpha
MMP: Matrix metalloproteinase
Mo1: Classical monocytes
Mo2: Intermediate monocytes
Mo3: Non-classical monocytes
MT: Myofibroblasic transdifferenciation
NCAVD: Non-calcific aortic valve disease patients
NF-κB: Nuclear factor-kappa B
OD: Osteogenic differenciation
OPG: Osteoprotegerin
OPN: Osteopontin
PBMCs: Peripheral blood mononuclear cells
PDC: Pyruvate dehydrogenase complex
PDK4: Pyruvate dehydrogenase kinase 4
RANK: Receptor activator of nuclear factor kappa B
RANKL: Receptor activator of nuclear factor kappa-B ligand
RNAseq: RNA sequencing
SAVR: Surgical aortic valve replacement
TAVR: Transcatheter aortic valve replacement
TIMP: Tissue inhibitor of metalloproteinases
TLR2: Toll-like receptor 2
TNC: Tenascin C
TNF: Tumor necrosis factor
VCAM-1: Vascular cell adhesion protein 1
VECs: Valvular endothelial cells
VICs: Valvular interstitial cells
Vol: Healthy volunteers

## 13. Supplemental Material

### Human peripheral blood collection

Between October 2021 and December 2023, the MIRACL prospective study enrolled 58 coronary artery disease patients without aortic valve disease (NCAVD) and 60 patients with severe CAVD at Rouen University Hospital. The study was approved by the CPP SUD-EST (RCB: 2021-A01850-41). Healthy volunteer (Vol) samples were obtained from fresh blood purchased from the French Blood Establishment (EFS) from 37 healthy anonymous donors, for whom no information on sex or age was available More details in Supplementary Methods.

### CD14^+^ monocyte isolation from human peripheral Blood and cell culture

CD14^+^ monocytes from NCAVD, CAVD, and Vol groups were directly isolated from peripheral blood by negative selection using EasySep^TM^ Human Monocyte Isolation Kits (StemCell Technologies, Cat#19669) according to the manufacturer’s instructions. Freshly isolated monocytes were either processed immediately for RNA extraction or cultured for 3 days in Dulbecco’s Modified Eagle’s Medium (DMEM, Gibco, Cat#41966-029) supplemented with 50 µM 2-mercaptoethanol (Gibco, Cat#21985-023), 10% foetal bovine serum (FBS, Gibco, Cat#10500-064), and 1% penicillin/streptomycin (Sigma-Aldrich, Cat#P4333). Cells were seeded at a density greater than 150,000 cells/cm² in 6-well plates (Corning, Cat#353046) for conditioned media collection.

### Human aortic valve collection and VIC cell culture

Between October 2021 and December 2023, calcified human aortic valve samples were obtained from 10 patients with CAVD undergoing surgical aortic valve replacement (SAVR). Patients included in the study had no history of infective endocarditis, bicuspid aortic valve, or rheumatic heart disease. The study was approved by the CPP NORD-OUEST I (RCB: 2016-A00137-44). Written informed consent was obtained from all patients prior to tissue collection. Find detailed protocol for human VICs isolation in Supplementary Methods.

### Flow cytometry

Peripheral blood mononuclear cells (PBMCs) were isolated from peripheral blood of NCAVD, CAVD patients, and Vol collected in EDTA tubes, by density gradient centrifugation using Ficoll. PBMCs were analysed by flow cytometry using an 18-color LSR Fortessa cytometer (BD Biosciences). Data analysis was performed with FlowJo software (TreeStar, Inc., San Carlos, CA). Results are expressed as % immune cell subpopulation per total cell count. More details in Supplementary Methods.

### RNA Extraction

Total RNA was extracted from at least 3.10⁶ sorted CD14^+^ monocytes and valvular interstitial cells (VICs) treated with monocyte-conditioned media, using the NucleoSpin RNA XS kit (Macherey-Nagel, Cat# 740902) according to the manufacturer’s instructions.

### RNA sequencing

CD14^+^ monocyte RNA samples extracted from volunteers (Vol, n=5), NCAVD (n=5), and CAVD (n=5) patients, with at least 400 ng of total RNA, with a concentration >20 ng/ml and a RIN ratio >8, were sent to Novogene Company Ltd (Cambridge, UK) for library preparation, mRNA sequencing, and bioinformatic analysis. The RNA-Seq data generated in this study have been deposited in the public Gene Expression Omnibus database under accession code GSE290104. More details in Supplementary Methods.

### RT-qPCR

A total of 1,500 ng of previously isolated total RNA was reverse transcribed using random primers (Invitrogen, Cat#48190-011) and M-MLV reverse transcriptase (Invitrogen, Cat#28025-013) according to the manufacturer’s instructions. cDNA synthesis was carried out at 37 °C for 1 hour, followed by 95 °C for 5 minutes, and a final hold at 4 °C. The cDNA was diluted 1:4 or 1:10 in nuclease-free water before amplification with LightCycler® 480 SYBR Green I Master (Roche, Cat#04-707-516-001) using the Rotor-Gene Q system (QIAGEN)Absolute gene abundance relative to the B2M housekeeping gene was calculated based on a cDNA standard curve using Rotor-Gene Q Series software version 2.3.5.1. Further details are provided in the Supplementary Methods.

### Quantification of VIC calcium production by O-Cresolphthalein complexone assay

VICs seeded at the density of 25,000 cells/cm² in 24-well plates were exposed for 10 days to conditioned media collected from CD14^+^ monocytes isolated from Vol, NCAVD, and CAVD patients and cultured as described above. Calcium deposition was directly quantified using the o-cresolphthalein complexone assay. Detailed protocols are provided in the Supplementary Methods.

### Multiplex Immunoassay

Conditioned media from circulating CD14^+^ monocytes of NCAVD (n=5), CAVD (n=5), and Vol (n=5) were analyzed using Bio-Plex Pro Human Inflammation Panel 1 (37-plex) and Human Chemokine Panel (40-plex) kits (Bio-Rad, Cat#171AL001M and Cat#171AK99MR2) on a Luminex xMAP bead-based multiplex system to quantify multiple analytes simultaneously.

### Immunocytochemistry

VICs, seeded at a density of 25,000 cells/cm² on 14 mm diameter coverslips in 24-well plates, were treated with monocyte conditioned media from NCAVD (n=5), CAVD (n=5), and healthy volunteers (Vol, n=5). After 24 hours, cells were stained with 20 µM OsteoSense 680 EX (PerkinElmer, Cat#NEV10020EX) to visualize calcium deposits, followed by fixation with 4% paraformaldehyde. Permeabilization was performed with 100% methanol, and cells were then blocked in PBS containing 1% BSA and 4% donkey serum. Subsequently, cells were incubated overnight at 4 °C with the respective primary antibodies. More details in Supplementary Methods.

### Statistical analysis

Data are expressed as mean ± SEM or percentages of control. Statistical significance was assessed using Kruskal–Wallis with Dunn’s post-hoc test or one-way ANOVA with Sidak’s multiple comparisons, depending on normality (Shapiro–Wilk test). Analyses were performed with GraphPad Prism 8. Differences were considered significant at p < 0.05; symbols in figures indicate ***p < 0.001, **p < 0.01, *p < 0.05, ns = not significant.

### 11. Supplemental Figures with Figures Legends

**Figure S1.**
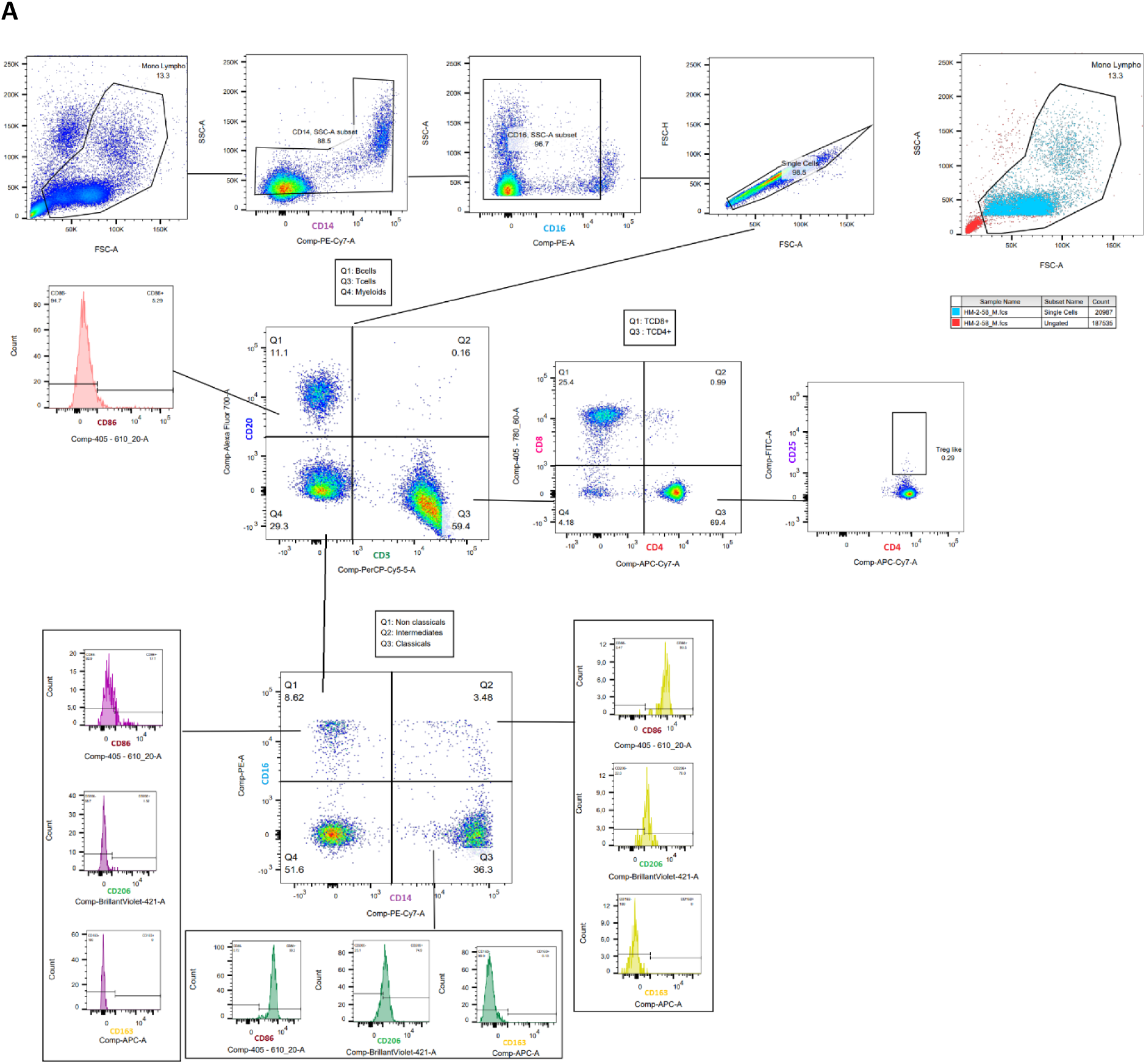
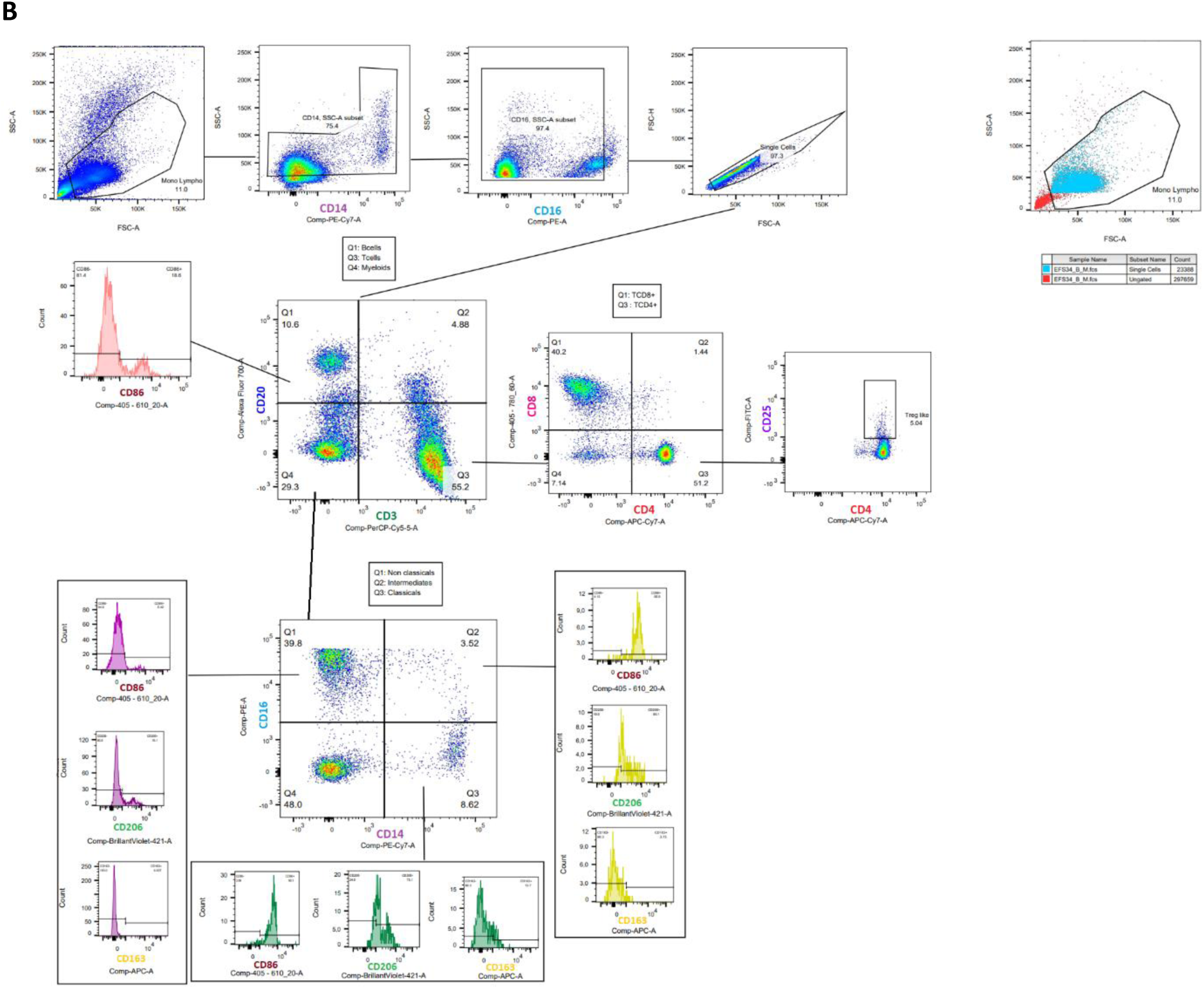
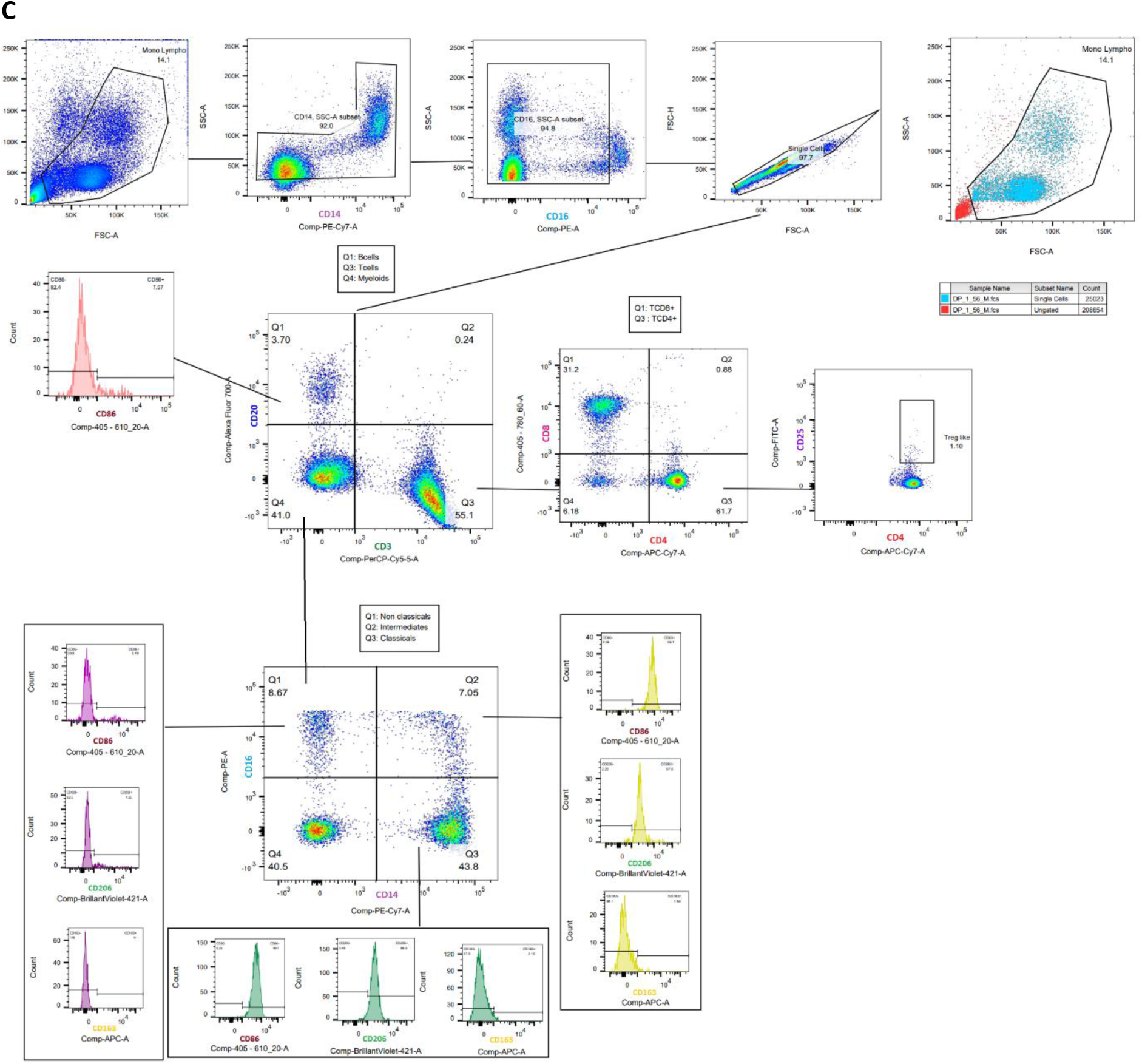
Flow cytometry gating strategy and staining panel for peripheral blood mononuclear cells (PBMCs) from (A) CAVD patients, (B) NCAVD patients, and (C) Vol. PBMCs were stained for an 11-color panel across 55 samples (NCAVD, n = 20; CAVD, n = 15; volunteers, n = 20). Each panel included lineage markers CD20, CD3, CD4, CD8, and CD25 to exclude B and T lymphocytes, and monocyte markers CD14 and CD16 to identify monocyte subsets. Additional markers CD86, CD163, and CD206 were used to characterize monocyte polarization toward unpolarized, pro-inflammatory, or immunomodulatory macrophage phenotypes, respectively. The gating strategy for identifying these populations is shown for each group.

**Figure S2:**
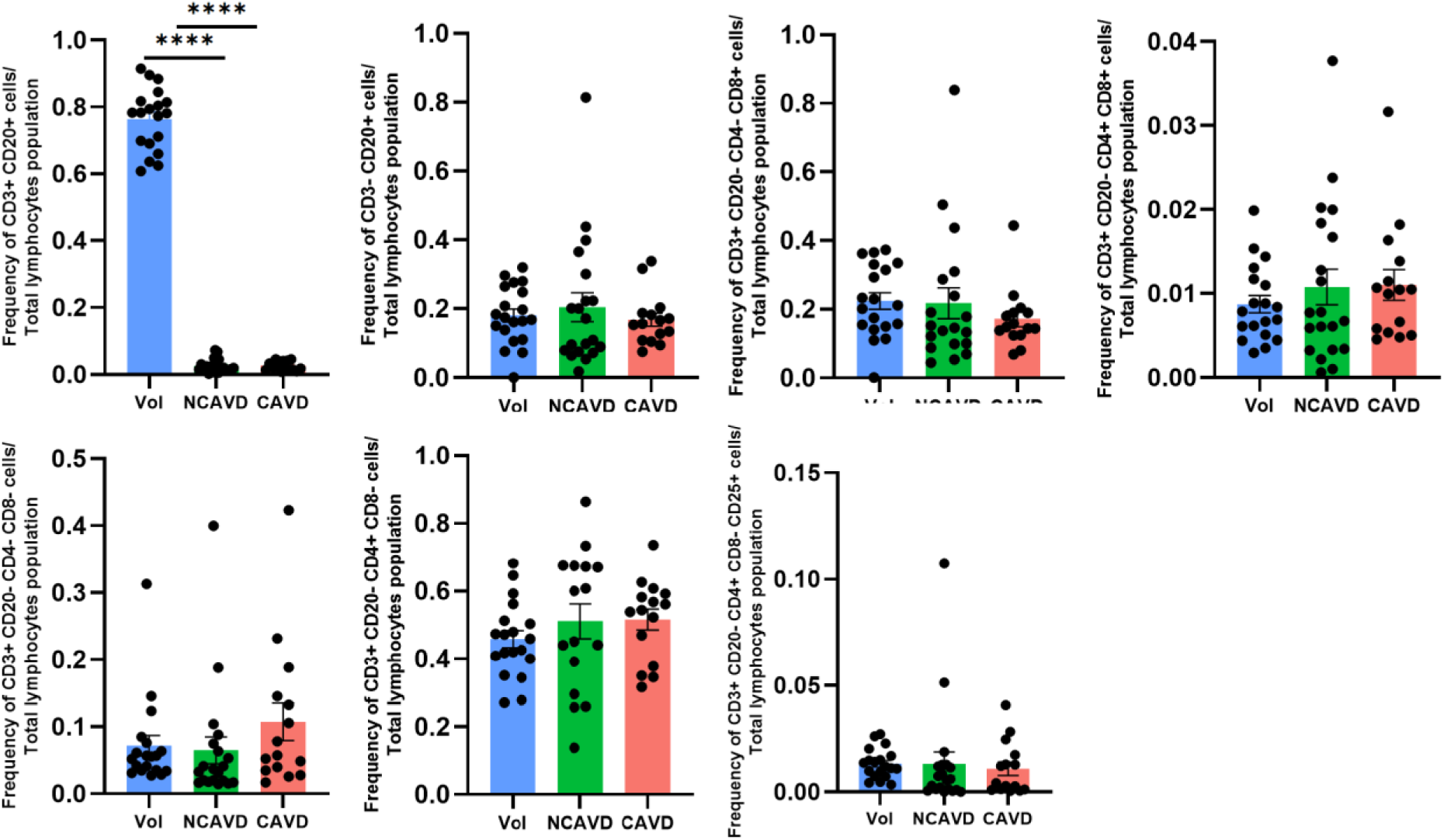
Lymphocyte’s subsets characterization in PBMCs from healthy volunteers, NCAVD and CAVD patients. After excluding unwanted events and doublets, T and B lymphocytes were identified using CD3 and CD20 markers. B lymphocytes were defined as CD3⁻CD20⁺, and T lymphocytes as CD3⁺CD20⁻. T lymphocyte subsets were further characterized using CD4, CD8, and CD25 markers. The frequency of each lymphocyte subpopulation was calculated as the ratio of the specific lymphocyte subset cell count to the total lymphocyte cell count. Data were analysed using one-way ANOVA followed by Sidak’s multiple comparison test. Statistical significance is indicated as follows: ****p < 0.0001.

**Figure S3:**
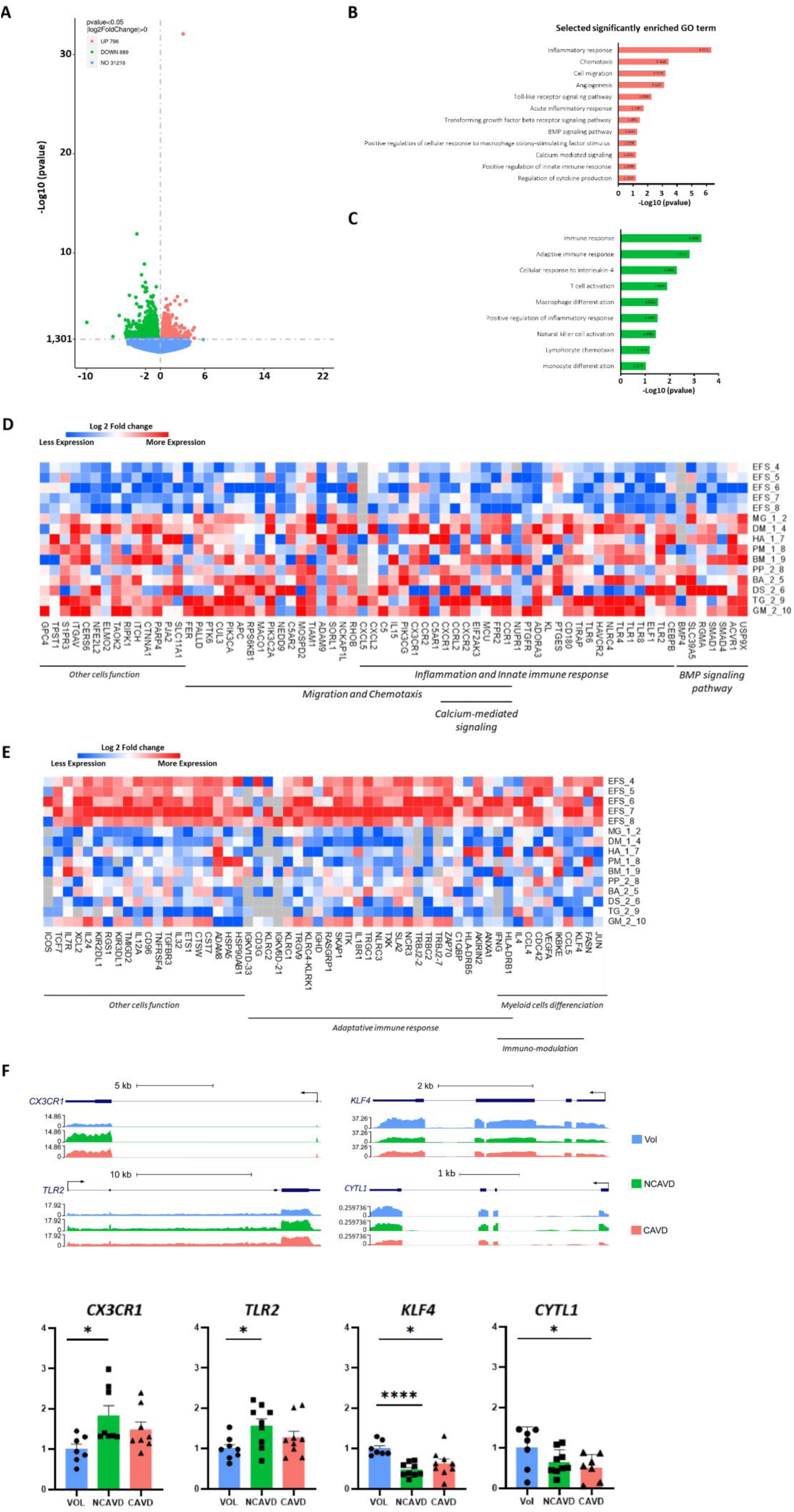
Transcriptomic analysis of dysregulated genes in CAVD CD14^+^CD16^-^monocytes. (A) Volcano plot showing upregulated (red) and downregulated (green) genes combining data from five donors in the healthy volunteer (Vol) group and five donors in the CAVD group. Genes were considered statistically significant with an adjusted p-value (padj) < 0.05, calculated using a negative binomial model (Novogene). (B, C) Bar graphs represent Gene Ontology (GO) biological processes enriched among upregulated (B) and downregulated (C) genes. (D, E) Heatmaps of selected significantly upregulated (D) and downregulated (E) genes identified by RNA-seq in human monocytes. (F) mRNA expression of genes of interest was measured by RT-qPCR and normalized to B2M. Sample sizes were as follows: for *CX3CR1, n=8* in the NCAVD and CAVD groups and n = 7 for the healthy volunteers (Vol) group; for *TLR2, n=9* in the NCAVD and CAVD groups and n = 8 for the healthy volunteers (Vol) group; for *KLF4, n=9* in the NCAVD and CAVD groups and n = 7 for the healthy volunteers (Vol) group; for *CYTL1, n=7* in the healthy volunteers (Vol) and CAVD groups and n = 9 for the NCAVD group. Statistical analysis was performed using one-way ANOVA followed by Sidak’s post-hoc test. Outliers were excluded. Statistical significance is indicated as *p < 0.05 and ****p < 0.0001.

**Figure S4:**
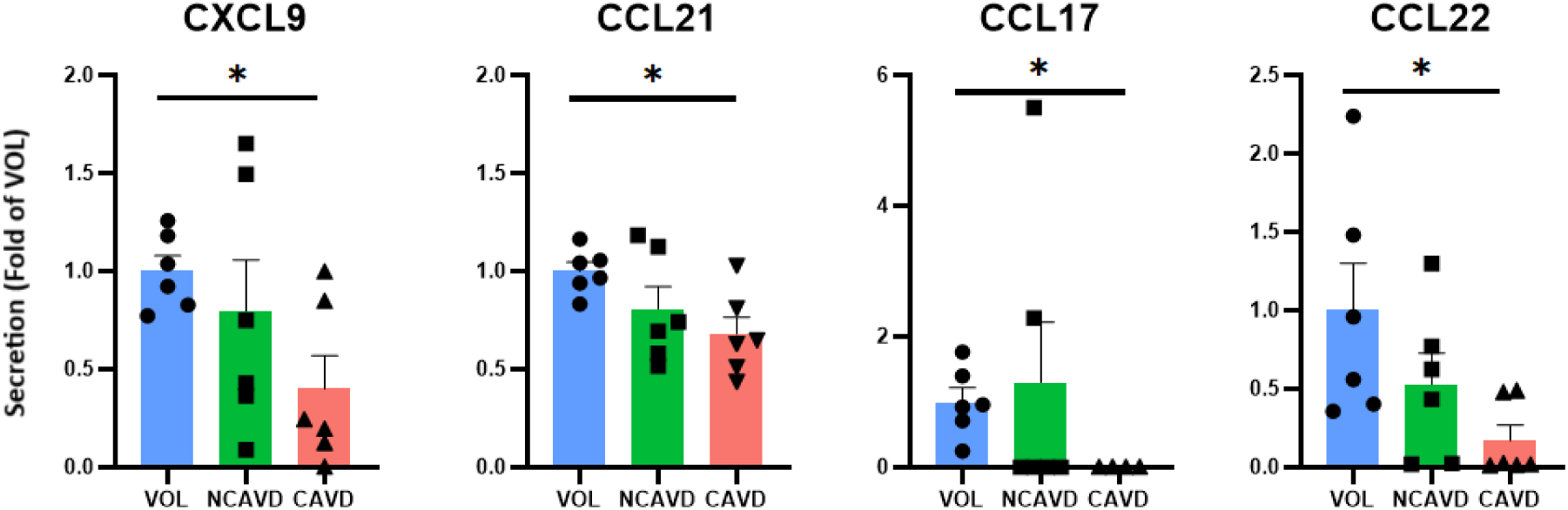
Secretions from CAVD CD14^+^monocytes exhibit reduced capacity to recruit T lymphocytes and to promote a Th2 immune response. The secretome of CAVD CD14^+^ monocytes is depleted in cytokines involved in T lymphocyte recruitment (CXCL9, CCL21) and Th2 response promotion (CCL17, CCL22). Data are normalized to the housekeeping gene B2M. Asterisks (*) indicate statistically significant differences compared to healthy volunteer samples (blue), determined by one-way ANOVA followed by Sidak’s post-hoc test. Outliers were removed. Statistical significance is denoted as *p < 0.05 and ***p < 0.001.

**Figure S5.**
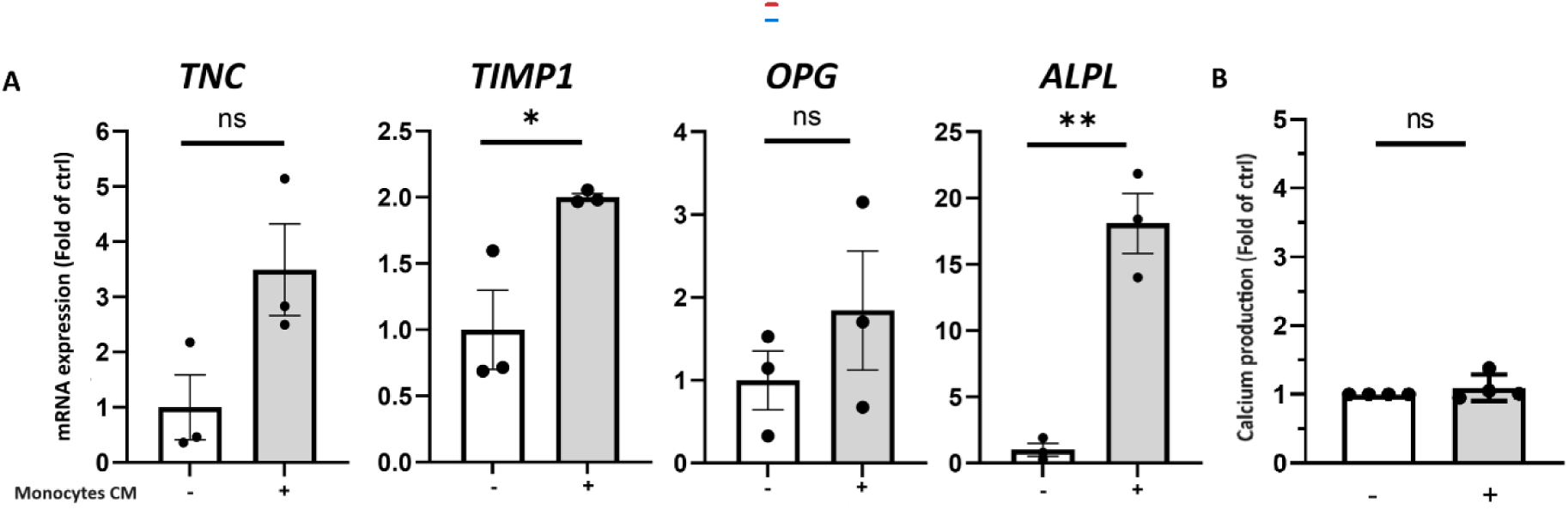
Conditioned medium from healthy donor CD14⁺ monocytes enhance VICS myofibroblastic transdifferentiation, has minimal impact on osteogenic transition, and does not increase calcium deposition. (A)Hydroxyapatite crystals were quantified using O-cresolphtalein assay (n=4 for each condition). *TNC, TIMP1, ALPL* and *OPG* gene expression in VICS (B) measured by RT-qPCR (n = 3 per condition) after treatment with CAVD monocytes conditioned media or non-conditioned media for 3 or 7 days. Gene expression is normalized to the housekeeping gene *B2M*. Statistical analysis was assessed with Mann-Whitney test. The p values are depicted as follows: * p<0.05. ns= non-significant.

